# Supervised and Unsupervised Classification of Cocoa Bean Origin and Processing using Liquid Chromatography-Mass Spectrometry

**DOI:** 10.1101/2020.02.09.940577

**Authors:** Santhust Kumar, Roy N. D’Souza, Britta Behrends, Marcello Corno, Matthias S. Ullrich, Nikolai Kuhnert, Marc-Thorsten Hütt

## Abstract

Liquid Chromatography-Mass Spectrometry (LC-MS) provides an unprecedented wealth of metabolomics information for food products, including insights into compositional changes during food processing. Here, we employed the largest available LC-MS dataset of around 300 cocoa bean samples to assess the capability of two popular multivariate classification methods, principal component analysis (PCA) and linear decomposition analysis (LDA), for studying bean geographic origin and responsible characteristic compounds.

The unsupervised method, PCA, only provides a limited separation in bean origin. Expectedly, the supervised method, LDA, provides a better origin clustering. However, it suffers from a strong, nonlinear dependence on the set of compounds used in the analysis. We show that for LDA a compound filtering criterion based on Gaussian intensity distributions dramatically enhances origin clustering of samples, thus increasing its predictive efficiency. In this form, the supervised method of LDA holds the possibility to identify potential markers of a specific origin.

## 1. Introduction

The classification of cocoa based on quality has been a major challenge in the chocolate industry, which is in industrial practice currently based on the rather simple cut-test, rather not reflecting cocoa’s vast chemical repertoire. Cocoa quality and geographic provenance define cocoa prices and quality. Assurance of cocoa authenticity protects manufacturers and consumers alike from commercial fraud.

Several of our research approaches have previously addressed classification of cocoa beans or cocoa products based on variety of cocoa constituent classes (D’Souza et al., 2017; Kumari et al., 2018; Megías-Pérez et al., 2018; Sirbu et al., 2018) using information-rich techniques based on actual molecular fingerprints (Aculey et al., 2010; Magagna et al., 2017; Vázquez-Ovando et al., 2015), or low-resolution techniques based on sum parameters (Guehi et al., 2010).

The used methods have been shown to not only assess the degree of fermentation of a cocoa samples, but also to discriminate between its origin based on its metabolome fingerprint (Acierno et al., 2016, 2018; Bindereif et al., 2019; D’Souza et al., 2017; Kumari et al., 2018; Marseglia et al., 2016; Oliveira et al., 2016). Metabolic differences of the cocoa metabolome are a consequence of distinct agricultural practice, climate and soil influences (Adeniyi et al., 2019; Arévalo-Hernández et al., 2019; Asare et al., 2017; Ehiakpor et al., 2016; Kongor et al., 2016, 2019) and are mainly based on genetic variability found in cocoa tree populations in different origin countries as illustrated by the group of Meinhardt on various occasions (Arevalo-Gardini et al., 2019; Gopaulchan et al., 2019; Lindo et al., 2018; Zhang and Motilal, 2016).

Liquid Chromatography-Mass Spectrometry (LC-MS) constitutes the most powerful technique for metabolomics analysis providing high resolution combined with high sensitivity. Cocoa products are among the most complex materials available to mankind with 10 000 to 35 000 individual peaks detectable in a single mass spectrum in processed cocoa (Kuhnert et al., 2013; Milev et al., 2014). When comparing a multitude of samples, powerful chemometric algorithms must be used to extract meaningful information from such datasets. At the same time, large sample numbers are challenging such algorithms to the extreme.

The state of the level of classification of origin of cocoa samples in the above-mentioned studies remained limited, because it is either at the level of continental regions or consider only few countries. Here we use a large data set comprising of 297 LC-MS profiles of aqueous methanolic extracts rich in polyphenolics and peptides (positive and negative ion modes) of unfermented or fermented cocoa beans, as well as selected cocoa liquors, sourced from various countries (10). The data has been gathered over a period of several years (2014-2018) and belongs to different stages of transformation in a typical cocoa processing pipeline. First, we demonstrate the limitations of the standard unsupervised go-to approach, principal component analysis (PCA) in clustering country of origin of cocoa samples. Second, we use a supervised approach, linear discriminant analysis (LDA), and show that its classification strongly depends on the number of features used in the analysis. We then outline a statistically motivated intuitive procedure for selecting features (compounds) and show that it greatly improves the outcome of LDA. We further show that this procedure of feature selection, which we refer to as Gaussian Feature Stability (GFS) requirement, also significantly improves prediction result for the country of origin achievable through LDA. The improvement in clustering of LDA can help in finding compounds which can differentiate between country of origin.

## 2. Materials and Methods

### 2.1 Details of LC-MS data

In the analysis we present here, a total of 297 LC-MS profiles of cocoa aqueous methanolic extracts rich in polyphenols and peptides was used. The LC-MS profiles are broadly categorized into two MS ion modes, three sample types and 10 origins (countries). These LC-MS profiles have been obtained over a period of five years (2014-2018). The actual details of sample preparation, extraction, standard protocols, carrying out the LC-MS experiment and data collection are given in our earlier published reports (D’Souza et al., 2017, 2018). The number of LC-MS profiles with respect to ion modes, sample types and their country of origin are provided in Table 1.

**Table 1.** Information about the LC-MS dataset. Country-wise and process-wise distribution of number of LC-MS samples in the dataset used in this study.

|  | Positive ion mode |  |  |  | Negative ion mode |  |  |  |
| --- | --- | --- | --- | --- | --- | --- | --- | --- |
|  | Unfermented | Fermented | Liquor | Sum | Unfermented | Fermented | Liquor | Sum |
| Brazil | 4 | 4 | 0 | 8 | 4 | 4 | 0 | 8 |
| Cameroon | 3 | 3 | 6 | 12 | 3 | 3 | 7 | 13 |
| Ecuador | 8 | 12 | 3 | 23 | 8 | 12 | 5 | 25 |
| Ghana | 0 | 0 | 5 | 5 | 0 | 0 | 5 | 5 |
| Indonesia | 14 | 16 | 0 | 30 | 14 | 16 | 0 | 30 |
| Ivory Coast | 16 | 16 | 9 | 41 | 16 | 16 | 9 | 41 |
| Madagascar | 0 | 0 | 0 | 0 | 0 | 0 | 5 | 5 |
| Malaysia | 6 | 3 | 0 | 9 | 6 | 3 | 0 | 9 |
| San Thome | 0 | 0 | 0 | 0 | 0 | 0 | 5 | 5 |
| Tanzania | 3 | 9 | 0 | 12 | 3 | 9 | 4 | 16 |
| Sum | 54 | 63 | 23 | 140 | 54 | 63 | 40 | 157 |

Positive and negative ion mode data jointly provide a more holistic approach enabling an instant molecular snapshot of the coca bean chemical composition. Negative ion mode data show a bias towards polyphenolic compounds, acids and carbohydrates, whereas positive ion mode reveal a multitude of peptidic structures. Both modes are largely complementary with an overlap of identified compounds of around 30-40 % depending on the sample type.

### 2.2 Data pre-processing and cleaning

We first processed the individual MS ion modes LC-MS data using MZMine (Pluskal et al., 2010). This yields a retention time aligned peak area list indicating the *m/z* ratio, the *retention time* and the integrated peak areas of each sample. Then using a precompiled list of compounds and their chemical formulas, corresponding *m/z* values were de-convoluted and assigned for determined in four ion types: [M-H], [M-2H], [M-3H], [2M-H], for negative ion mode, and [M+H], [M+2H], [M+3H], [2M+H], for positive ion mode. Using this information, the compounds detected in the LC-MS data were assigned to known structures, whenever possible based on authentic references, tandem MS data or literature precedents, else the compound was included, however, considered as ‘Unknown_’ suffixed with the *m/z* value (e.g. Unknown_865.1927). In cases where more than one compound matched an *m/z* value, a combined name, joined by ‘or’ was assigned. The assignment procedure was performed for obtaining further insights, when possible, about the detected compounds. These processed data were then combined with metadata about each sample in a single data structure, which contained information about the sample type, origin and peak areas of various compounds. The sum of peak area values belonging to each sample was normalized to 100, so the peak area in the sample represent relative percentage amount of compound in the LC-MS profile of the sample referred to the sum of all intensities (percentage normalized peak area). Henceforth, we refer to the percentage normalized peak area as the peak areas itself. Columns of compounds were sorted in descending order by their mean peak area across all samples, such that the compounds with higher mean peak areas are placed in earlier columns, and those with lesser peak areas are placed in later columns.

### 2.3 Unsupervised and supervised learning methods

A couple of multivariate statistical analysis or ‘machine learning’ methods, both unsupervised (e.g., PCA – Principal Component Analysis) and supervised (e.g., LDA – Linear Discriminant Analysis, Random Forests, Support Vector Machines, Neural Nets) have been used in this study. These methods were applied using the popular scikit-learn module (Pedregosa et al., 2011) of Python programming language. Standard scaling was implemented while performing PCA. In case of Random Forests, Support Vector Machines, and Neural Nets, where it is possible to vary algorithm specific parameters, default parameter settings in the implementation in scikit-learn were used, to avoid additional complexity of dealing with parameter specific algorithmic performance.

### 2.4 Gaussian Feature Stability criterion

The Gaussian Feature Stability (GFS) criterion is based on the test whether a set of values is distributed normally, or not. If the values in the peak area list corresponding to a compound in the LC-MS profiles, under a given sample type and belonging to a given origin, are found to be normally distributed, we say that the compound satisfies the Gaussian Feature Stability requirement. For testing normality of a set of values, we use the Shapiro-Wilk (Shapiro and Wilk, 1965) test implemented in the popular Scipy module (Jones et al., 2001) of Python programming language. We test normality at a p-value threshold of 0.05.

### 2.5 Null model

First, we find out how many compounds (say *n*) satisfy GFS criterion under a group of samples. Here, a group is comprised of samples belonging to a given country (say *X*), under a given sample-type (say *Y*). Then we randomly choose *n* compounds from the same group of origin (*X*) and sample type (*Y*). Using these *n* randomly chosen compounds, we calculate prediction from LDA. This procedure is repeated a number of times to obtain result statistics. The result so obtained is referred to as result from null model.

## 3. Results and Discussion

For this study we have selected around 300 samples representing well the world of industrial cocoa, including high production origins from Africa, South America and Asia over several years of harvest. The dataset includes as well cocoa beans of different processing stages reflecting chemical changes along the processing chain. In wine chemistry for example a memory effect with respect to origin was recently proposed allowing improved origin prediction following ageing (Roullier-Gall et al., 2014). In the main text we report results, primarily, about the negative ion mode data. When appropriate or needed, corresponding plots, results, etc., for the positive ion mode are reported in the Supporting Information.

### 3.1. Classification of cocoa using PCA

Principal component analysis (PCA), an unsupervised method of classification (James et al., 2013), has been the first choice “go-to tool” in food chemistry analyses for exploring and studying grouping relationships in samples (Aculey et al., 2010; Cordella, 2012; Granato et al., 2018). However, PCA has been found of limited success in the study of country of origin of cocoa beans (D’Souza et al., 2017; Kumari et al., 2018). In Figure 1, we show PCA score plots for the negative ion mode data using unfermented, fermented and liquor samples of cocoa in our analysis (using top 2000 compounds; see Supporting Information for positive mode data and plots with different numbers of compounds). It can be seen that samples belonging to same country (depicted by dots with same color in the PCA plots) tend to be present close by. However, one can also clearly witness the mixing of samples belonging to different countries. We note that this separation or mixing of samples further varies with (a) the number, and (b) the actual set, of compounds used as features in the analysis. This points to a need for an alternative approach for identifying marker-compounds, which can potentially allow distinction between cocoa samples belonging to different country of origin.

**Figure 1.**
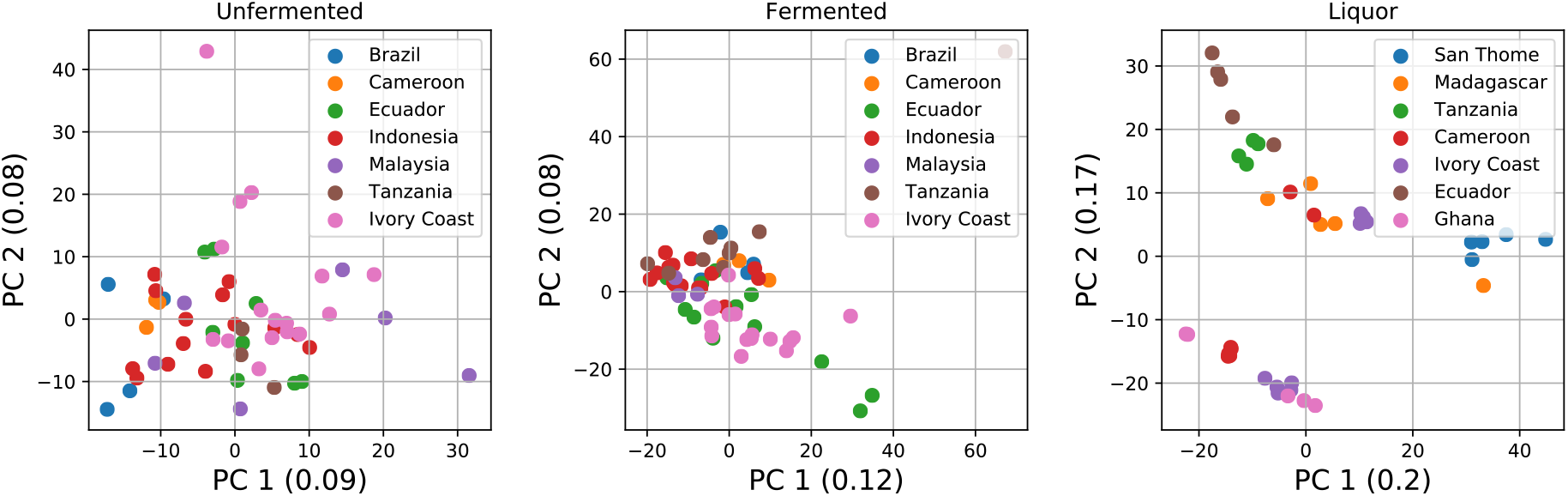
PCA of Unfermented, Fermented and Liquor samples obtained from Cocoa processing pipeline. The PCA gives a limited amount of grouping of samples belonging to same country.

### 3.2. Classification of cocoa using LDA

We next applied linear discriminant analysis (LDA) (James et al., 2013) to the cocoa samples analyzed by LC-MS. LDA, as a supervised method of classification, uses the available class information of the samples (in our case, country of origin) in order to find out axes, which give best possible grouping of samples belonging to the same class (i.e., country, in our case). It achieves this by simultaneously minimizing the within-class variation and maximizing between-class variation. This is in contrast with PCA, which determines axes with most variation independent of the class the samples. In this way, LDA is better suited for determining compounds, which are good differentiators of samples belonging to different origins and compounds, which makes samples belonging to the same country similar.

In Figure 2, we show the LDA of unfermented, fermented, and liquor samples. For each sample type, the analysis is performed under four divisions, using: (a) all compounds, (b) the first 500 most abundant compounds, (c) the first 100 most abundant compounds, and (d) only the first 10 most abundant compounds. It becomes apparent that the grouping of samples varies with number of abundant compounds (or features) included prior to performing the LDA. A marked improvement in the grouping of samples belonging to same origin country is observed when only including the top 30-50 most abundant compounds for the LDA. However, this soon disappears as the number of compounds are further decreased.

**Figure 2.**
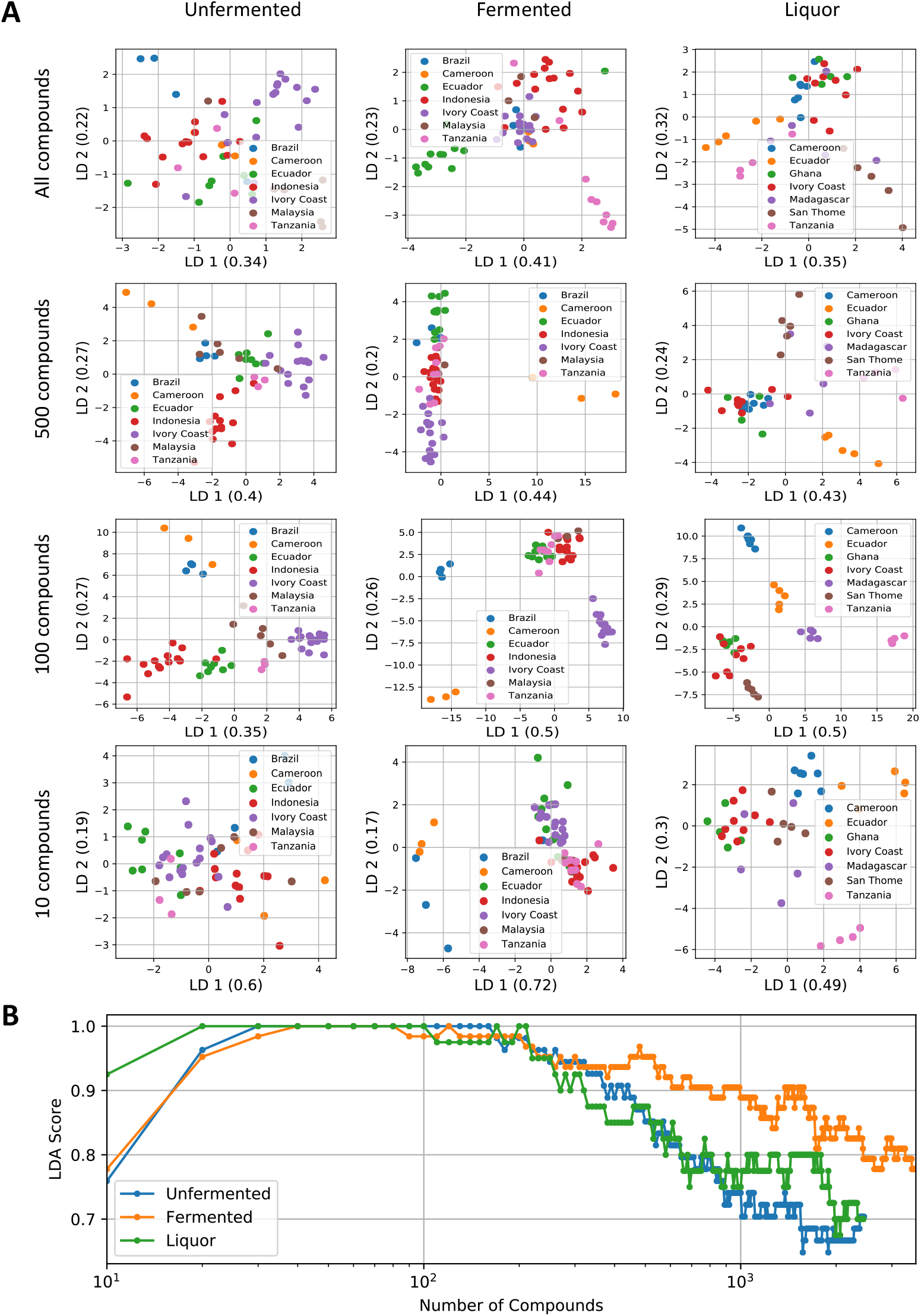
LDA for different sample-types using different number of compounds (features). **(A)** The scatter plots show that the quality of grouping of samples depends on the set of compounds used in the analysis. **(B)** Non-linear variation of LDA-score (an estimate of clustering of same samples) as a function of number of compounds employed in the LDA.

The quality of grouping of samples of same type can quantified by calculating the LDA score. While in originality the LDA score is a measure of prediction power of the trained LDA model upon the test dataset, it works well as a measure of clustering of the datapoints (or the *classifiability* of the data), when the test dataset is kept the same as the training dataset. In Figure 2B, we show the variation of LDA score with the number of compounds used for the analysis. The classifiability from LDA changes with the number of compounds in a non-linear manner. In the Supporting information, we provide results on positive ion mode and the corresponding plots with the number of compounds displayed on a linear (not logarithmic) scale. This points to the criticality of the number and set of compounds used as features in performing the LDA, and how to select them.

### 3.3 Filtering data for relevant features

In general, feature selection is a factor that needs to be decided before performing a multivariate analysis. Before performing LDA upon a given sample-type, we find out the set of compounds which follow a Gaussian distribution under each country under the given sample type. We refer to this filter as Gaussian Feature Stability (GFS) requirement (see section 2. Materials and Methods). It is fair to expect that the distribution of intensities for a compound belonging to a given country under a given process category is centered on some stable value, which might be characteristic for this subset of samples. Otherwise, if the amount of the compound in the samples belonging to a given country under a process category is not stable enough, it cannot be considered as a reliable feature enabling classification for the samples. The reason for instability can be varied: ranging from error in faithful detection of the compounds to the influence of other factors such as subsamples being procured in different batches over a period of time (as in our case the data is gathered over a period of few years). After obtaining such a list of compounds adhering to the GFS criterion, we use them as features for performing the LDA. Figure 3 shows the resulting improvement in the LDA brought about through this approach. Essentially, the GFS criterion is able to reduce conflicting information from samples belonging to same country.

**Figure 3.**
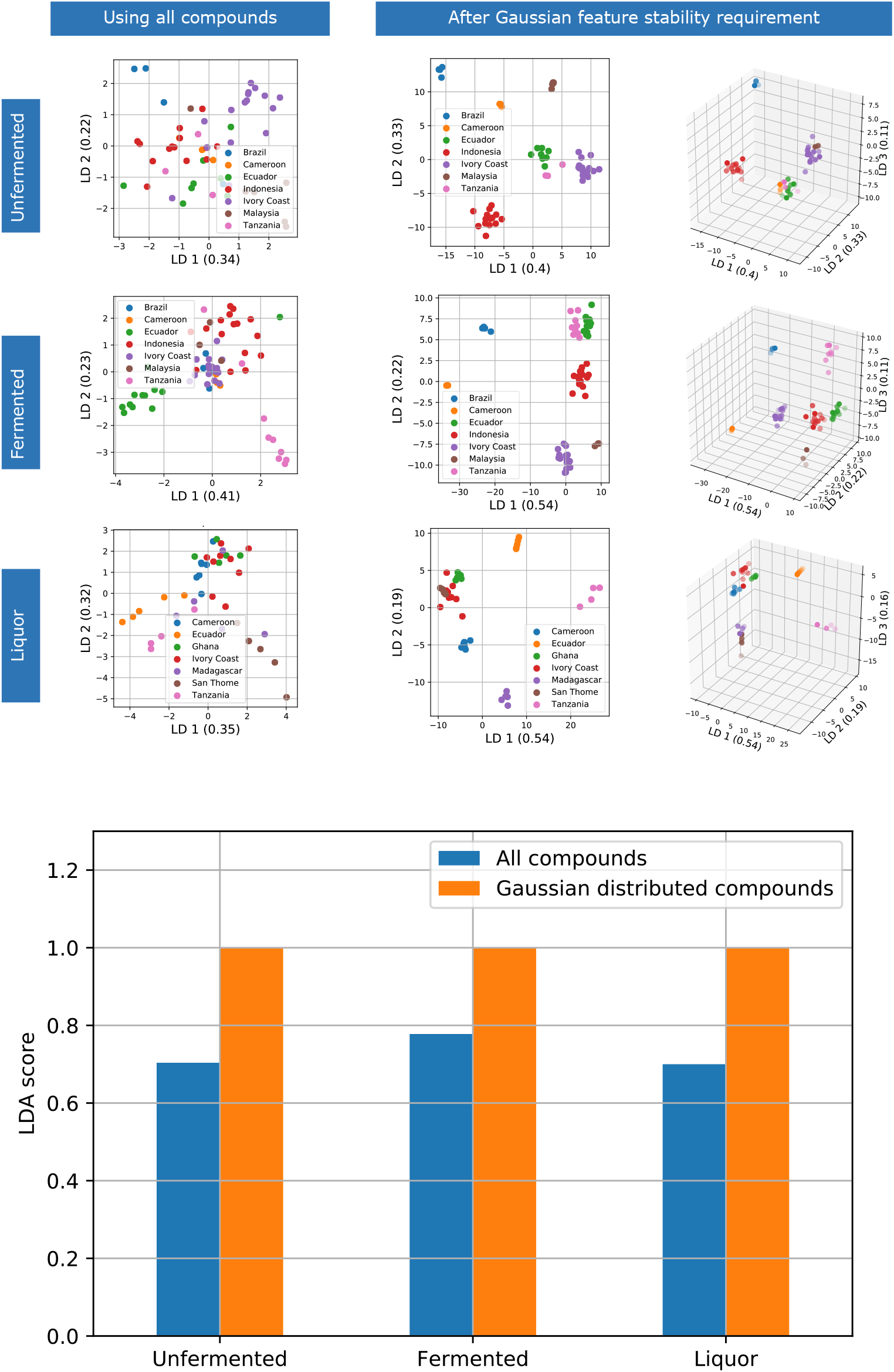
LDA clustering improvement after Gaussian Feature Stability (GFS) requirement. **Scatter plots:** Visual representation of clustering in LDA before and after application of GFS. **Bar chart:** Quantitative comparison of improvement in clustering in LDA after application of GFS.

It should be pointed out that once the country clusters become clearly separated by the LDA, the LDA score used above will no longer discriminate the amount of cluster separation (giving a value close to 1 for all cases). The higher quality of the reduction of compounds based on GFS will become clear in the following, when the LDA result is used for predictive purposes. The origin-wise distributions of peak-list area of some metabolites and peptides obtained after applying GFS requirement for unfermented, fermented, and liquors, are shown through boxplots in Figure 4. The compounds have differing country-wise distribution, for e.g., *p-Coumaroyl aspartate [M-H]* serves as marker within the liquor category. Identifying compounds with stable and differing distributions can be helpful in identification of origin specific characteristic compounds. Figure 4 shows some selected examples of relative quantities of marker compounds within the three stages of cocoa processing investigated.

**Figure 4.**
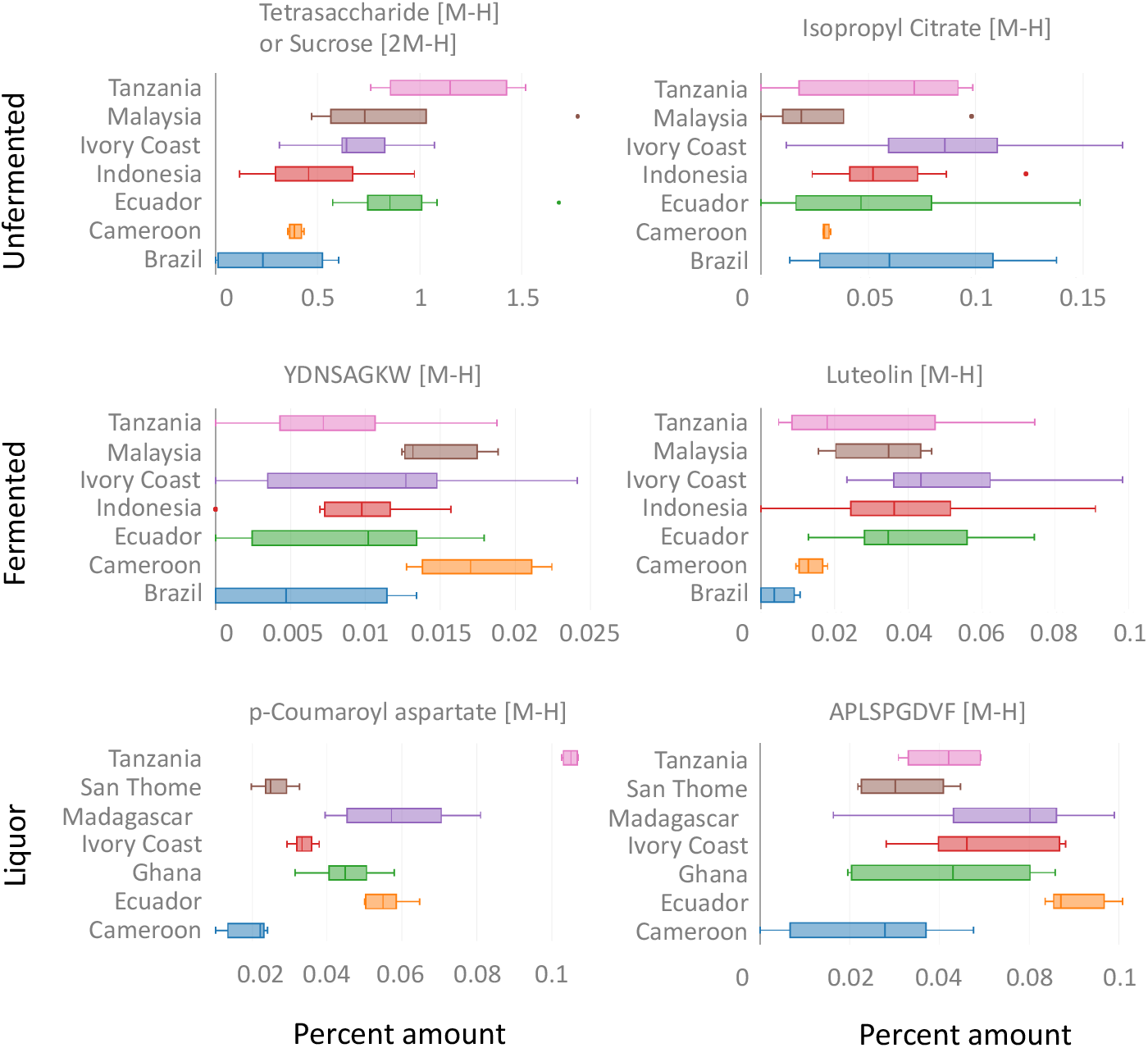
Profiles of compound concentration: The figure shows profiles of variation of concentration of compounds belonging to different countries for some compounds obtained after the application of Gaussian Feature Stability requirement. It can be seen that the countries have differing distributions of concentration of compounds.

### 3.4 Effect of Gaussian Feature Stability on prediction through LDA

Next, we study the effect of Gaussian Feature Stability (GFS) criterion on *prediction* of country of origin through LDA. For each sample-type category, the list of compounds satisfying GFS criterion was found. The sample dataset belonging to each category was divided into training and test sets in ratio 3:1. After that, the LDA model was trained through the training dataset, and its predictive power was assessed through test dataset by calculating the LDA score. The whole procedure, from data splitting to prediction score, was repeated several times (100) to achieve result statistics. To put the outcome into perspective, a comparison was made with an appropriate null model. In this null model, the same number of compounds as obtained previously by GFS criterion, was randomly chosen from all compounds under a given category and used to obtain the prediction statistics (see 2. Materials and Methods). This procedure was repeated 100 times and was done for each category.

The result is plotted in Figure 5, with blue bars representing prediction power of LDA, when compounds satisfying GFS requirement were used, and orange bars represent the prediction power, when same number of randomly chosen compounds were used. It can be seen that higher predictive ability of LDA was achieved, when the compounds chosen through GFS criterion were used, as compared to the case when same number of randomly chosen compounds were used (2. Materials and Methods, Null model). Prediction using compounds satisfying GFS requirement through some popular machine learning algorithms is given in Supporting Information for comparison with that of LDA.

**Figure 5.**
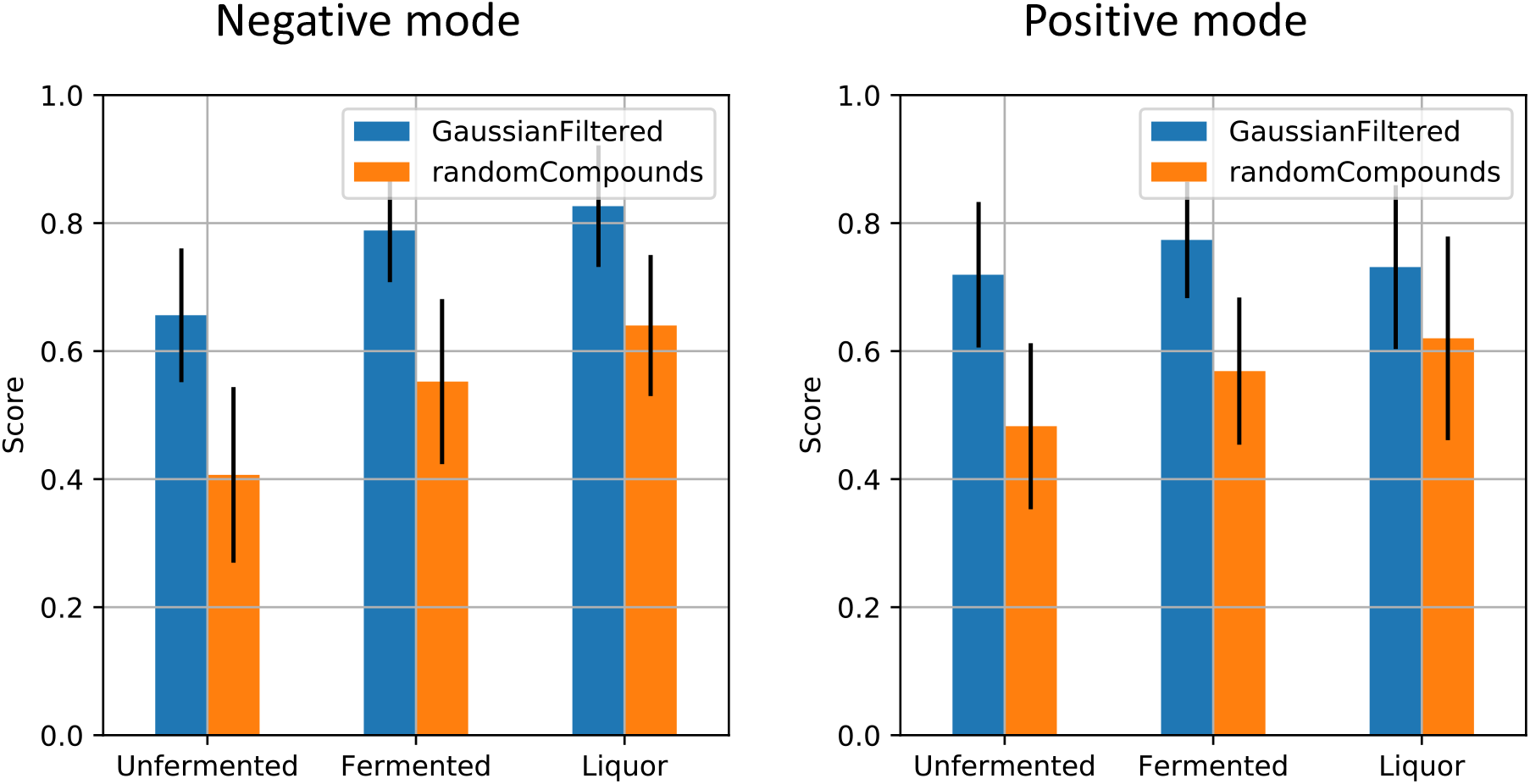
Improvement in LDA predictive capability upon usage of compounds satisfying GFS requirement: Using the set of compounds obtained from GFS to predict country of origin gives higher predictive power (blue bars) than expected by chance (orange bars). The chance predictability was calculated by taking same number of compounds as obtained by GFS requirement, but chosen randomly, from each sample type (unfermented, fermented and liquor) from the list of available compounds in respective positive/negative ion mode datasets.

## 4. Discussion and Conclusions

We have used the biggest dataset of the LC-MS chemical profile of cocoa available to date to study the feasibility of bean origin classification. Data from both positive and negative ion modes have been analyzed.

PCA has often been used as a standard tool for studying such cases. PCA projects the samples in the dataset into a lower dimensional space whose axes are formed by the principal components, which are vectors obtained from the linear combination of the original features, such that the variation is maximized along the principal components. The samples in the space thus obtained show most variation present in the dataset. The amount of contribution of the original feature (in our case, compound) in the determination of the principal axes can be read from a corresponding loading plot and is a measure of the importance of that feature (compound) in obtaining the represented variation in the dataset. If there appear groups of samples in the obtained space, then the compounds having high contribution in the determination of principal axes are designated as compounds responsible for separating the dataset into the obtained groups. There is quite some possibility that the compounds, which have most variation, might be able to separate the dataset into some groups; however, it is not necessary. Only when the groups obtained after PCA show some correspondence to the original groups in the dataset, can one say with some confidence that compounds with relatively higher weights in the principal axes of the space determined by PCA are responsible for the differentiating one group from another. In summary, generally speaking, there is no binding reason for the compounds which show more variation in the dataset to also be the ones which differentiate one group of samples from another in the dataset. While PCA has been shown to successfully separate beans on basis of their fermentation status, it fails to separate beans on the basis of their country of origin. Thus, there is a need to move beyond PCA to identify compounds, which can differentiate beans of one country from another.

Since we have the country information about samples in our dataset, we employ LDA, which is a supervised learning method. Like PCA, LDA gives a lower dimensional space whose axes are represented by the obtained linear discriminants and in which samples in the dataset can be visualized. However, unlike PCA, LDA uses the class information about the samples in the data set to provide a space in which the within class variation is minimized and between class variation is maximized. In our case, this means that the samples belonging to the same country should be as closely placed together as possible, and in turn these groups simultaneously should be placed as far as possible. Again, like the PCA, the weighted contributions of the original feature axes in the determination of linear discriminants determine the importance of features in the observed separation, if any. Here we illustrate that LDA can be used to obtain a good grouping of samples belonging to the same country, however, this depends on the subset and number of compounds used while performing LDA. The set of compounds is thus somewhat arbitrary. Further, a large number of low intensity compounds gets dropped out of analysis some of which might still be relevant differentiators of origin of beans.

We coupled LDA with a statistical procedure of data cleaning which is intuitive and realistic: We ask which of the compounds among the given set of compounds has a Gaussian distribution (or nearly so) for each of the countries they are present in. One may argue that the characteristic features of a group with have their values distributed around some mean value along with some random fluctuations, thus the distribution should be near Gaussian. This criterion on one hand finds out statistically stable features, and on the other hand removes noise in the data. We then perform LDA with the compounds, which pass this criterion that we call Gaussian Feature Stability requirement. We notice a marked improvement in LDA classification of the samples in the datasets.

Thus, this work makes an advancement in the direction of finding plausible differentiator compounds for the origin of beans. Further, we couple a supervised learning method with a novel, intuitive and realistically motivated criterion of feature selection and noise reduction.

## Acknowledgements

We thank Nina Böttcher excellent technical support in sample logistics and preparation. This work was funded by the COMETA project, which is financially supported by Barry Callebaut AG. Barry Callebaut also provided samples for analysis. It has released the article for publication.

