## Supporting Information for "Supervised and Unsupervised Classification of Cocoa Bean Origin and Processing using Liquid Chromatography-Mass Spectrometry"

### 1 Classification using PCA

#### 1.1 PCA—positive mode (2000 compounds)

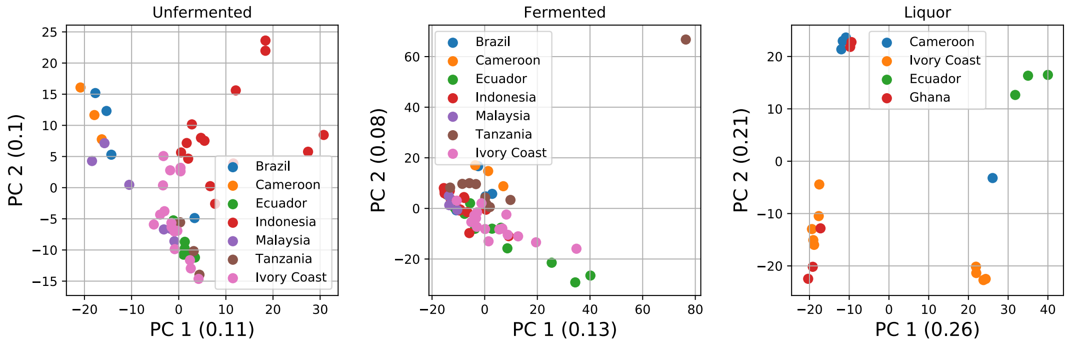

#### 1.2 PCA—varying number of compounds (negative mode)

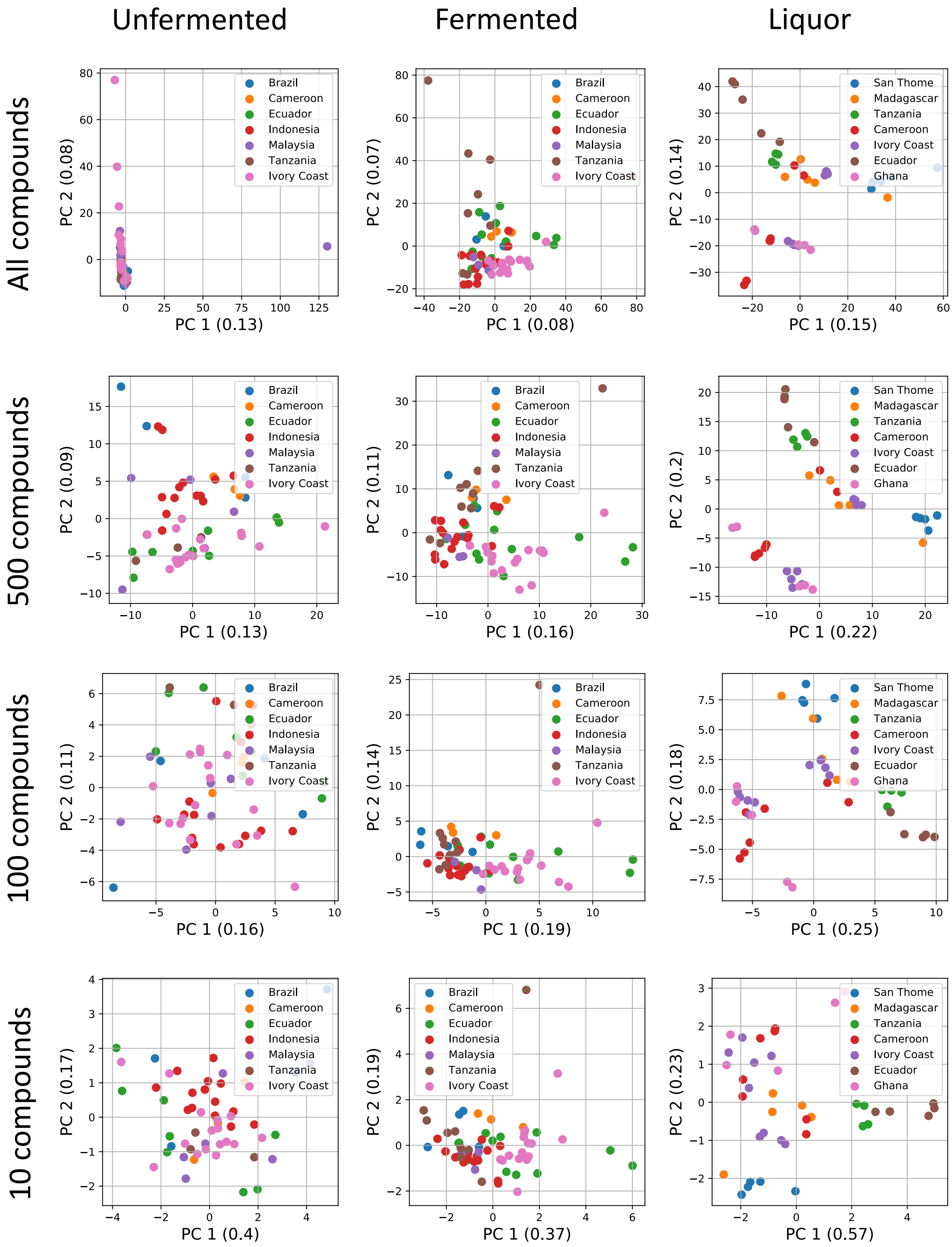

#### 1.3 PCA—varying number of compounds (positive mode)

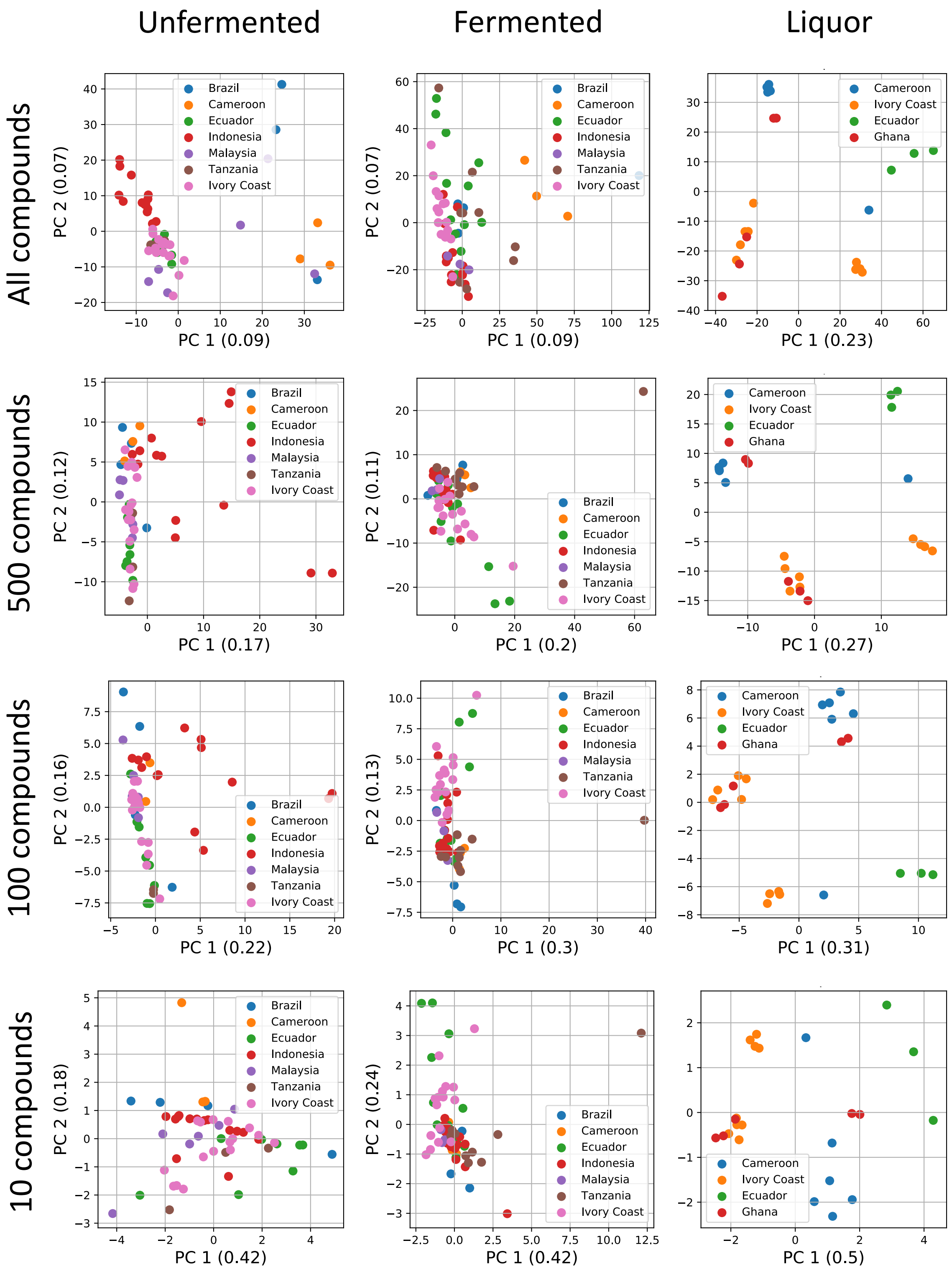

### 2 Classification using LDA

#### 2.1 Clustering variation with different number of compounds—positive mode

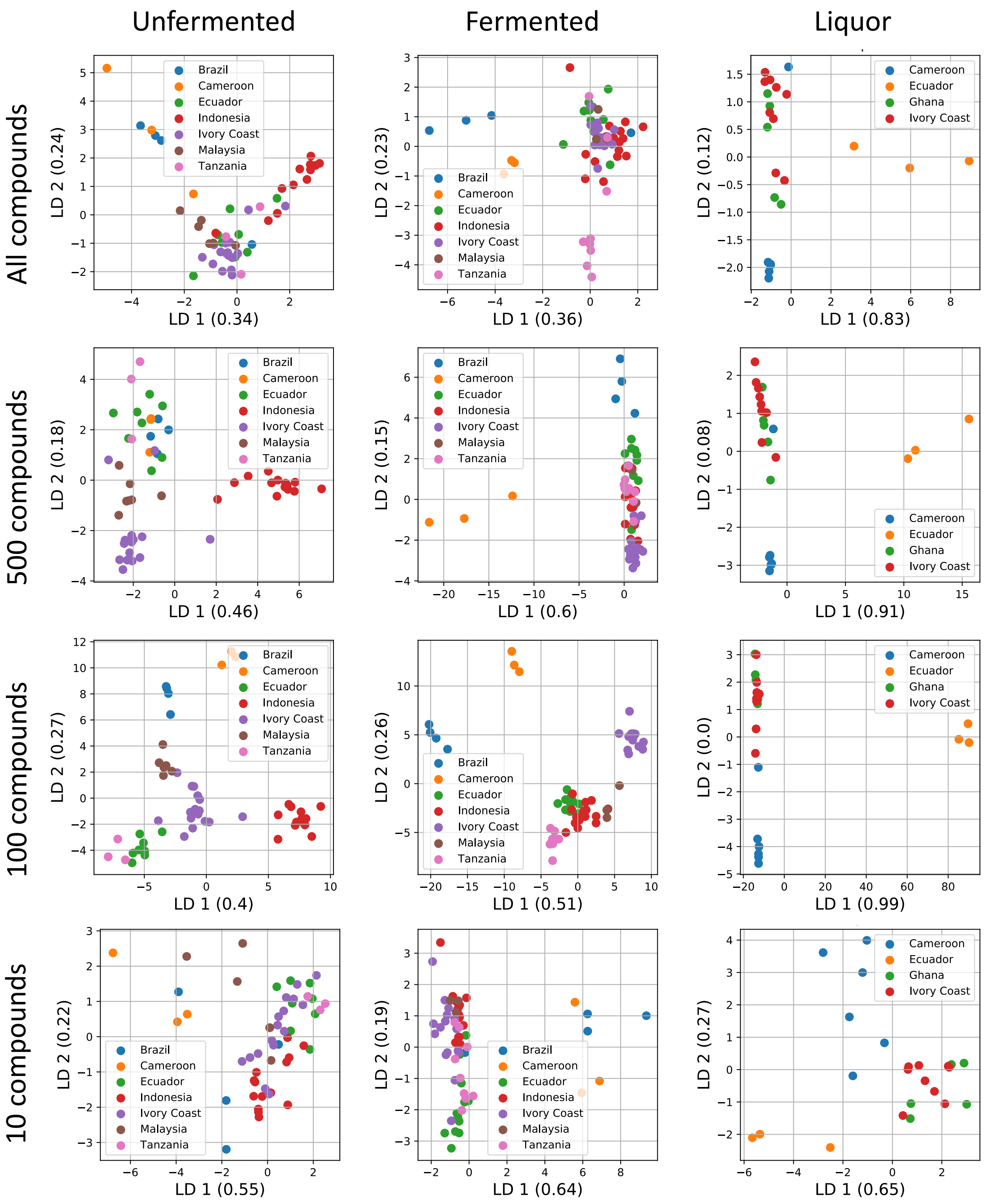

#### 2.2 LDA-score variation—negative mode

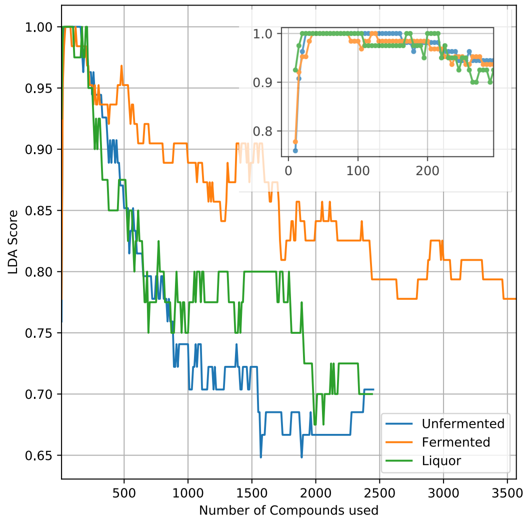

#### 2.3 LDA-score variation—positive mode

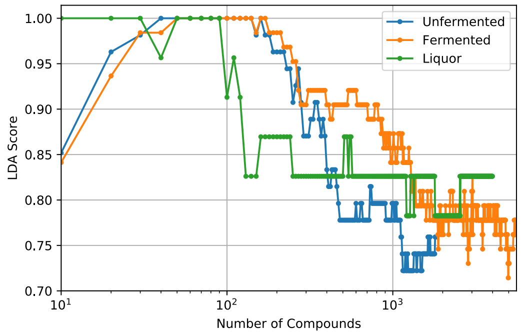

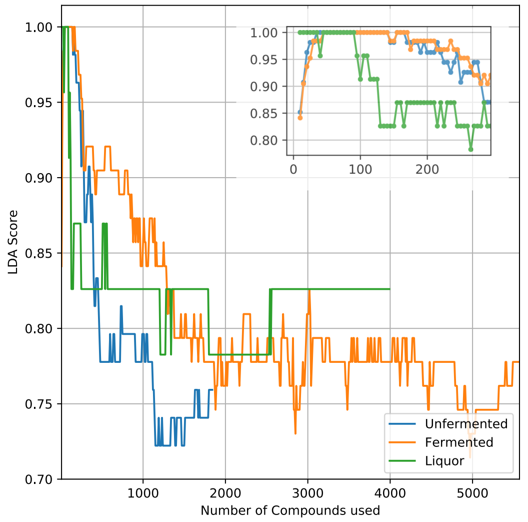

### 3 Filtering data for relevant features (Gaussian filtering within countries)

#### 3.1 LDA clustering improvement after GFS—positive mode

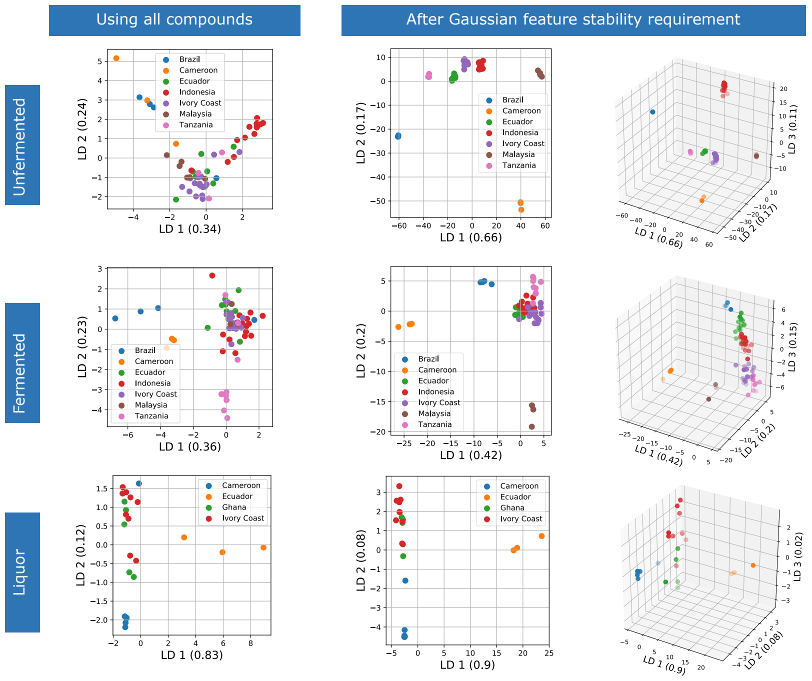

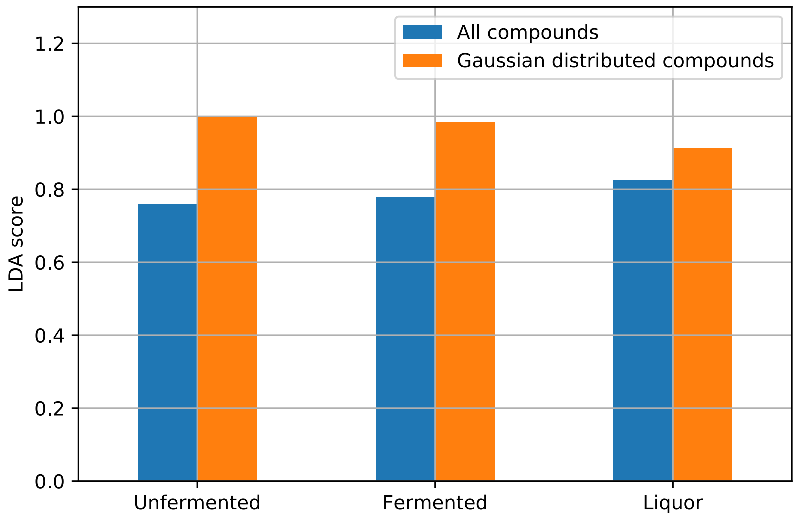

#### 3.2 Profile of concentration of some compounds—positive mode

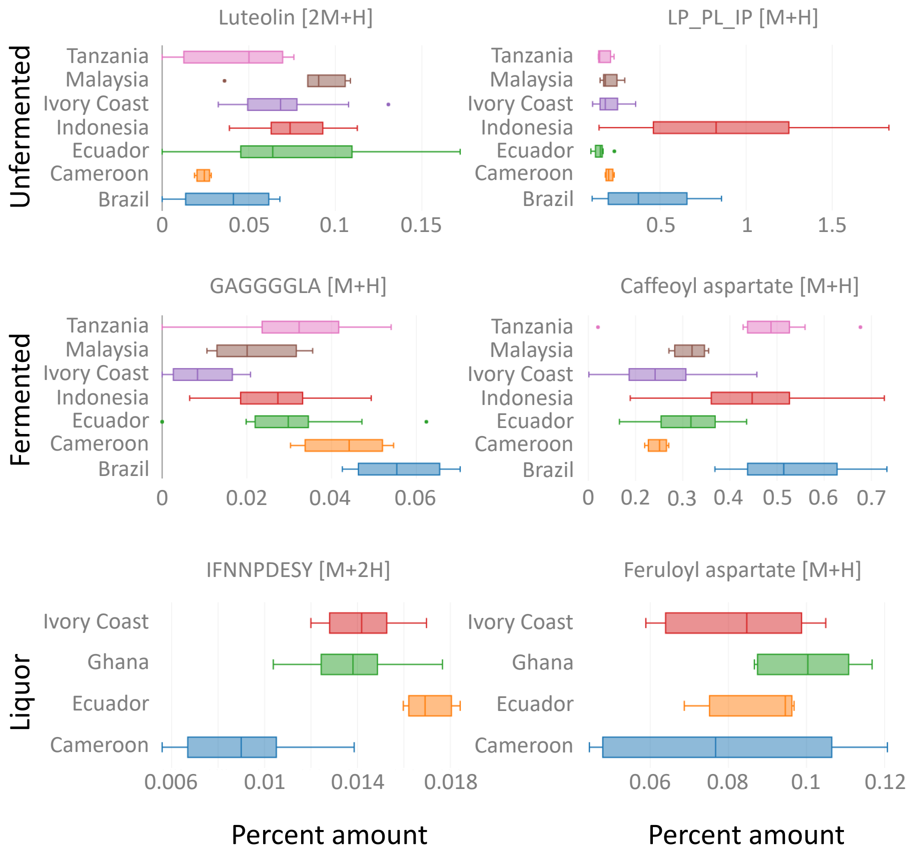

### 4 Prediction through some popular ML algorithms using compounds satisfying GFS

#### 4.1 Result from negative mode data

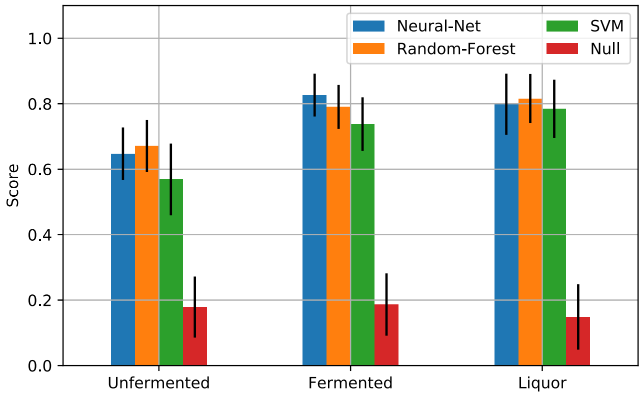

#### 4.2 Results from positive mode data

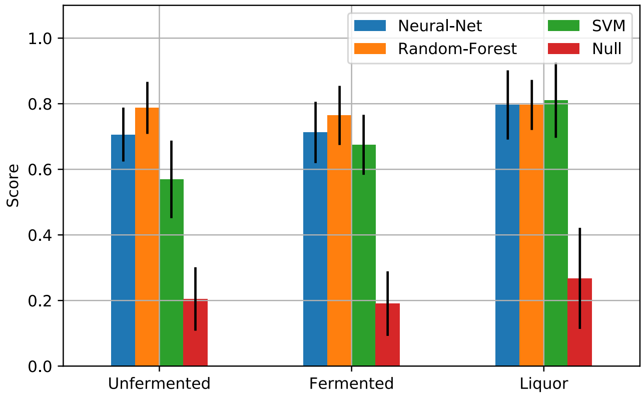
